# Semaphorin 3C attracts MGE-derived cortical interneurons in the deep migratory stream guiding them into the developing neocortex

**DOI:** 10.1101/2021.03.18.435991

**Authors:** Kiara Aiello, Jürgen Bolz

## Abstract

While it is known that Semaphorin 3C acts as a guidance cue for axons during brain development, their potential role during interneuron migration is largely unknown. One striking observation is that Sema3C demarcates the pallial/subpallial border and the intracortical pathway of cortical interneurons in the dorsal telencephalon. Moreover, migrating cortical interneurons express Neuropilin1 and Neuropilin2, described receptors for Semaphorin 3A, 3F and 3C. All these reasons prompt us to examine possible roles for Sema3C on cortical interneuron migration.

Using several *in vitro* approaches, we showed that Nrp1-expressing MGE-derived interneurons from the deep migratory stream migrate towards the increasing Sema3C gradients. In contrast, inhibitory neurons from the superficial migratory stream that express Nrp2, do not respond to this guidance cue. In the present study, we proposed that diffusible Sema3C expressed in the Pallium provides a permissive corridor that attracts the Nrp1-expressing interneurons from the DMS into the dorsal telencephalon.

## 1. INTRODUCTION

Compromising only 20-30 % of the neurons in the cortex, inhibitory cortical neurons play a key role in processing and storing information [1]. Their positioning and integration during brain development are crucial for accurate brain function, the misplacement of cortical interneurons is linked to neurological or neuropsychiatric disorders like autism, epilepsy, anxiety, and schizophrenia [2–4].

One fascinating aspect of cortical interneurons is that they originate far away from their final position, migrating long distances from the basal to the dorsal telencephalon. During migration, interneurons follow distinct paths. Each path depends from which embryonic eminence of the basal telencephalon interneurons are derived and when. Depending on the time and domain of origin, interneurons will express different transcription factors, which translate in a particular set of receptors and ligands confining them to each path. In the mouse, around embryonic day 11.5 (E11.5) early-born interneurons originate from the medial ganglionic eminence (MGE) and the preoptic area (POA). These cells predominantly migrate marginally along the superficial migratory stream (SMS), and enter the cortex in the marginal zone (MZ). In contrast, from E15.5 to E16.5, later-born interneurons, predominantly from the MGE and lateral ganglionic eminence (LGE), follow the deep migratory stream (DMS) along the ventricular / subventricular zone (VZ/SVZ). From E12.5 to E14.5, the MGE seems to be the main source of cortical interneurons that migrate either through the deep or superficial migratory stream. [5–12]

Several route-specific membrane-bound and secreted factors are described to orchestrate the tangential migration and to channel the distinct migratory streams into the appropriate routes, most of them are already known as brain wiring molecules that navigate axons. Cortical inhibitory neurons avoid repellent molecules, and move towards increasing gradients of attracting cues. Currently, most of the described chemokines are repellent and the only known attractive factor is Neuregulin-1, which is recognized by MGE-derived interneurons that express the corresponding receptor ErbB4. The membrane-bound isoform of Neuroregulin-1, expressed in the basal telencephalon, represents a short-range cue. Dissimilarly, the diffusible isoform in the cortex is secreted out of specific cells in the surrounding environment, and has been suggested to act as a long-range cue [14–16].

Another group of secreted guidance molecules is Class-III Semaphorins (Sema3). Sema3 and some of their co-receptors Neuropilins, are widely expressed in the developing brain and were already described to navigate axons during brain development. Additionally, they are known to play an important role during the migration of inhibitory neurons by mediating repulsive activity and proliferation [6, 7]. The developing striatum expresses Sema3A and 3F that prevent Nrp-expressing cortical interneurons from entering this non-target area. In contrast, striatal neurons downregulate their Nrp levels, which allows them to enter the striatum. [6, 7, 18]

Sema3C is also a wiring molecule from the Class-III Sema family. The interaction of Sema3C and its co-receptors Nrp1, Nrp2 has been shown to attract developing cortical axons [20, 21]. This wiring molecule is expressed in the cortical SVZ and IMZ, where the interneurons from the DMS enter the dorsal telencephalon. At the same time, co-receptors of Sema3C, Nrp1 and Nrp2, are expressed complementary by migrating interneurons en route to the dorsal telencephalon. In the deep migratory stream interneurons mostly expressed Nrp1 and in the superficial migratory stream interneurons expressed mostly Nrp2 receptor [5, 6, 9]. However, the effect of Sema3C on migrating interneurons is not known. This raises the question, whether this molecule influences the migratory behavior of inhibitory cortical neurons.

In the present study, we examined the possible role of Sema3C as a guidance molecule for migrating cortical interneurons. Using several *in vitro* approaches we showed that Nrp1-expressing MGE-derived interneurons from the DMS migrate towards increasing Sema3Cgradients. In contrast, inhibitory neurons from the SMS that express Nrp2 do not respond to Sema3C.

## 2. MATERIAL AND METHODS

### 2.1. Animals

Animals Time pregnant mice of the Nrp1 knockout strain [18] and Nrp2-deficient mice [19] were used. Also, wildtype mice from the C57BL/6 and NOR strains. The day of insemination was considered as embryonic day 1 (E1). Mice were bred and maintained under standard conditions with a 12 h light / dark cycle and access to food and water *ad libitum*. All animal procedures were performed in accordance with institutional regulations of the Friedrich-Schiller University Jena (Germany).

### 2.2. Cell lines

We used stable Human Embryonic Kidney cell lines (HEK-293) expressing recombinant Sema3A or Sema3C tagged to an Alkaline Phosphatase (-AP) and resistant to Geneticin [19]. After thawing and the first passage, cells were kept in culture medium with 0.5 % Geneticin. Untransfected HEK cells were used as control.

### 2.3. Tissue preparation

Time pregnant mice were deeply anesthetized using peritoneal injection of 10 % chloralhydrate. The uterus was removed and embryos were dissected.

#### 2.3.1. Dissociated and explant MGE-neurons

Embryonal dissected brains were kept in Gey’s balanced salt solution (GBSS) supplemented with 0.65 % D-glucose. For the binding, stripe and boyden assay, MGE was dissected completely after opening each hemisphere exposing the dorsal and ventral telencephalon. Subdomains of the basal telencephalon were collected in ice-cold Hank’s balanced salt solution (HBSS, Invitrogen) supplemented with 0.65 % D-glucose (HBSS/Glucose). MGE tissue was incubated in HBSS with 0.25 % trypsin in bather bath (17 min at 37 °C). Afterwards, the tissue was dissociated by trituration and filtered through nylon gauze to remove cell aggregates. In the case of explant preparation for the cu-culture assay, the isolated MGEs were shopped in ice-cold collection medium (HBSS/Glucose) using a Tissue Chopper (McIlwain, 250 μm slice) or using a scalpel. The explants obtained were incubated in a methylcellulose medium in order to get round (1 h at 37 °C).

#### 2.3.2. Coronal brain sections

Embryonic heads were fixed in 4 % PFA (in 1X PBS, pH 7.4) overnight at 4 °C, followed by a sequential Sucrose treatment (10 %, 15 %, 30 % in 1X PBS, pH 7.4) as cryoprotection. After freezing in Isopentan and dry ice at – 40 °C, the heads were sectioned into 20 μm slices (at −21°C) using a cryotome CM1900 (Leica) and mounted on Superfrost Plus slides (Thermo Fisher Scientific).

### 2.4. Immunostaining

Single cells were fixed at 4 % PFA (in 1X PBS, pH 7.4) for 10 min at room temperature (RT). Brain slices and dissociated single cells were washed in 1X PBS (pH 7.4) with 0.2 % Tween 20, followed by blocking (in 4 % BSA in 1X PBS/0.2 % Tween 20) for 2h at RT and incubation with the primary antibody at 4 °C, overnight. After washing, the secondary antibody was applied for 2h at RT. The nuclei were stained with DAPI (1 ng/ml in H2O) for 15 min at RT. As primary antibodies were used: goat anti Nrp1 (R&D, 1:100), goat anti Nrp2 (R&D, 1:100), rabbit anti Calbindin (Swann, 1:600) and Placental alkaline phosphatase (PLAP) (ABD-serotec, 1:100). As secondary antibodies were used: donkey anti rabbit FP 647H (Interchim, 1:1000) and donkey anti goat Alexa555 (Molecular probes, 1:1000). In the case of the co-localization, after fixation, one additional washing step was performed with warm PBS (65 °C) to eliminate any intrinsic alkaline phosphatase present in the cells. At the stripe assay, in order to visualize the Human Sema3C- Fc recombinant chimera, bound to the stripes, anti-human conjugated to Alexa 488 (Invitrogen, 30 μg/ml), was applied after fixation, following the same procedure as with a secondary antibody.

### 2.5. *In situ* Hybridisation

For Sema3C and Nrp1, coronal sections of E14 and E16 embryos were obtained and treated as described in Bagnard et al., [17] and Ruediger et al., [16].

### 2.6. Binding assay

To test the binding of recombinant Semaphorins to MGE-derived interneurons, 100 μl dissociated neurons were plated with a density of 80.000 cells/100 μl culture media onto Laminin/PLL HNO3-glass coverslip, (19.5 μg/ml Laminin (Sigma-Aldrich) and 5 μg/ml Poly-L-Lysine (PLL, Invitrogen)). Warm FBS-containing culture medium was used for 2 h (Dulbecco’s Modified Eagle Medium (DMEM, Invitrogen) supplemented with 10% FBS, 10000 U/ml penicillin, 10000 μg/ml streptomycin, 0.065 % D-glucose and 0.4 mM Lglutamine). Next, the culture medium was replaced for neurobasal serum-free culture medium (37 °C).

Cells were incubated for a total of 46 h and exposed for 2 h to control or Sema3C-AP tenfold conditioned media diluted in freshly warmed FBS-free neurobasal medium (37 °C). Control conditioned medium or Sema3C-AP conditioned media were obtained as already described [22, 23]. The dissociated cells were fixed for 15 min with 4 % PFA/PBS and further immunostaining were performed.

### 2.7. Stripe assay

The stripe assay was performed as recently described by Rudolph et al., [22] using silicone matrices obtained from the Max-Planck Institute for Developmental Biology (Tübingen, Germany). We injected in each matrix 25 μg of Recombinant Sema3C-Fc (Sigma-Aldrich) in a concentration of 50 μg/ml in PBS. After incubation at 37°C for 1 h, HNO3-coverslips were washed with PBS and coated with laminin/PLL (for 30 min at 37 °C). 100 μl dissociated neurons were seeded at a density of 90.000 cells/ 100 μl and incubated in a humid atmosphere in the culture medium for 2 days at 37 °C and 5 % CO2. Then fixation (4 % PFA/PBS at RT) and immunostaining was performed.

### 2.8. Transwell migration assay

The Boyden chamber assay was performed as previously described by Steinecke at al., [25]. We used the haptotaxis kit Cell Biolabs (8 μm pore size, collagen-pre-coated). 500 μl of neurobasal FBS-free culture medium containing tenfold control or Sema3C-AP conditioned media was placed in the lower wells of the Boyden chambers. Conditioned media were obtained as already described [22, 23]. Next, we added in the upper compartment 300 μl single cell suspension (90.000 cells/100 μl media). After 6 h *in vitro*, cells were fixed (4 % PFA/PBS).

### 2.9. Coculture assay

As previously described by Ruediger et al., [18], HEK cell aggregates were placed in the middle of a drop of 20 μl chicken plasma (Sigma-Aldrich) on non-coated HNO3-cover slips. 5 to 7 explants were placed in the chicken plasma with a distance of 1 mm from the HEK cells. Next, we added 20 μl of fresh GBSS/Thrombin mixture (10.000 U thrombin, (Sigma) stored in 7,94 mL bidistillate water. From this 30 μl were resuspended to a total of 1ml GBSS,). After coagulation for 15 min at RT, 2 ml culture medium was added. Cocultures were incubated in a humid atmosphere for 2 div at 37 °C and 5 % CO2. The cocultures were fixed (30 min, 4 % PFA/PBS) and embedded in Mowiol. This procedure was performed with untransfected, Sema3A-AP or Sema3C-AP HEK cells aggregates.

### 2.10. Detection and analysis

Pictures of *in situ* hybridization experiments and the following *in vitro* assays were taken using a Zeiss Axiovert S100 inverted microscope (Zeiss, Germany) in combination with a digital camera (Spot, Diagnostic instruments). For the stripe assay, analysis was performed using ImageJ. Significance was calculated, applying a paired student’s t-test. Three independent preparations were performed (out of three litters). For the boyden chamber assay, analysis was performed using a manual counter blindly.

Statistical differences were determined using a two-tailed student’s t test. Results from each experimental condition are from two independent preparations (out of two litters).

Pictures from co-localization assays and immunolabelling of brain slices were performed with a confocal laser scanning TCS SP5 (Leica) and the software Leica Application Suite Advanced Fluorescent lite (LAS Af lite; Leica 2011).

Regarding the coculture assay, analysis were performed using ImageJ, only explants that outgrowth in all their surfaces were taken into account and the guidance index was scored as described by Marin et al., [12] and Flames, Long et al., [16]:

#### 2.10.1. Guidance Index

“0” was given if the explant had equally outgrowth; −1 or −2 for moderate or strong repulsion, when cells are migrating away from the explant and +1 or +2 for moderate or strong attraction when cells are migrating towards the explant. The guidance index was calculated from the averages of all scores from control, Sema3A-AP or Sema3C-AP cocultures.

#### 2.10.2. Migration distance

The distance migrated by MGE-derived neurons form VZ/SVZ explants, was measured on micrographs of each explant. First, the initial shape of the explant was digitally divided in six parts with one division parallel to the border of the HEK aggregate. On this way, it was determined a sector proximal and distal to the HEK aggregate [12, 16, 19]. As illustrated in Figure 3F, in each proximal and distal area, only the 10 farthest neurons out of the explant were taken into account. Second, the migrated distance was measured using the ImageJ program by making a line between the initial border of the MGE explant and the soma of the cell. Results for each experimental condition are from at least three independent preparations (out of at least three litters).

## 3. RESULTS

### 3.1. Sema3C -Nrp1 interactions in MGE-derived neurons

Several class-III-Semaphorins are expressed in the basal telencephalon and have already been described to repel migrating cortical interneurons from entering non-target areas like the striatum [5, 6, 7]. In this study, we detected the class-III-Semaphorin 3C in the developing telencephalon, which potential function on interneuron migration has not been described yet.

We performed *in situ* hybridisation on coronal brain sections at E14, using a specific ribo probe against Sema3C and its best characterized coreceptors Nrp1 and Nrp2. We detected a spatially distinct expression in the upper SVZ and IMZ of the dorsal telencephalon, where the interneurons migrating through the DMS enters the developing cortex. We further recognized a weak signal in the MGE along the SMS. Later at E16, there is a superposition of Sema3C and Nrp1 mRNA signal in the SVZ of the cortex (Fig. 1A).

**Figure 1.**
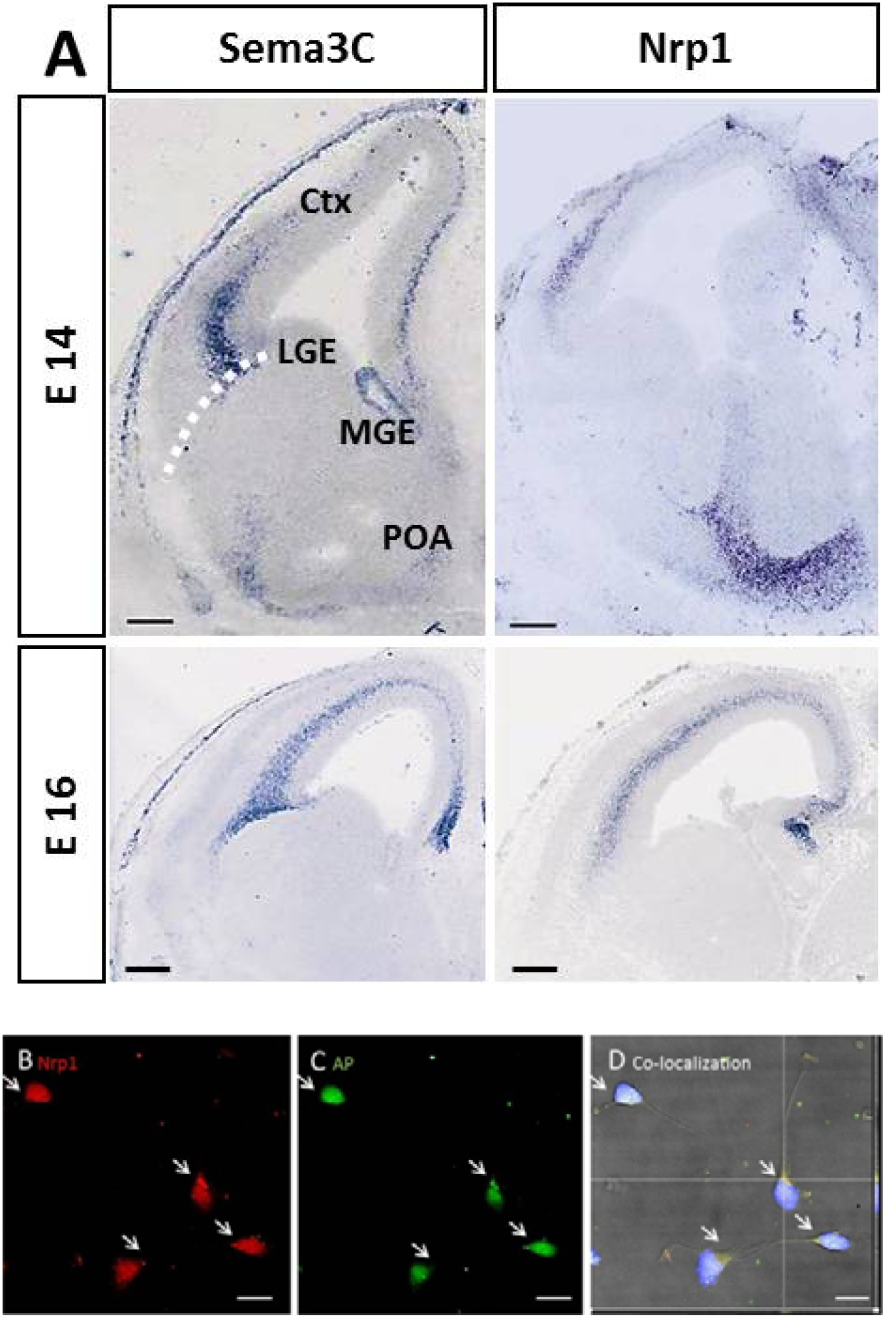
Sema3C expression pattern in the telencephalon and colocalization with Nrp1 receptors on MGE-derived single cells. (A) *In situ* hybridization on coronal sections revealed an expression of Sema3C in the SVZ and IMZ in the dorsal telencephalon at E14 and E16. The white dotted line illustrates the pallial/subpallial border. Scale bars equals 250μm. (B-D) E14.5 single neurons from the MGE were stimulated 2 h with Sema3C-AP or control conditioned medium. After fixation, double immunostaining was performed against (B) Nrp1 in red and (C) AP in green. (D) Co-localization of AP tag fused to Sema3C and Nrp1 is illustrated in an X and Y line scan, through a single optical plane. White arrows highlight AP/Nrp1 co-localization. Two independent experiments were performed. Scale bars equals 10 μm. E14.5, Embryonic day 14; Nrp1, Neuropilin 1; AP, Alkaline phosphatase. Ctx, Cortex; IMZ, intermediate zone; SVZ, Subventricular zone; MGE, Medial ganglionic eminence; LGE, Lateral ganglionic eminence; POA, Preoptic area; E, Embryonic day.

Therefore migrating MGE-derived interneurons expressing Nrp1 could interact with the Sema3C present in the SVZ of the cortex. To confirm this hypothesis, binding assays were performed. Thus, MGE-derived cortical single cells at E14.5 were prepared and incubated with recombinant Sema3C-AP or Control, respectively. Afterwards the AP was visualized using a specific antibody in combination with labelling against Nrp1 (Fig. 1B-C). The binding assay demonstrates that Sema3C can indeed bind to Nrp1, a guidance receptor that is present on cortical interneurons and therefore potentially influence tangential migration.

The complementary expression pattern of the coreceptors Nrp1 and the guidance molecule Sema3C prompt us to study an attractive effect of Sema3C on migrating Nrp1- MGE-derived interneurons.

### 3.2. Semaphorin3C, a permissive cue for MGE-derived inhibitory cortical neurons

To generally test if Sema3C has a chemotactic effect, we first performed a classical choice assay on alternating protein stripes, as previously described [24]. Thereby alternating stripes of recombinant Sema3C-Fc or control were coated on coverslips and dissociated MGEderived cells were homogeneously seeded on the protein stripes. The motile cells can freely choose their environment of preference. After 2 div, the cells were fixed and the cell number per stripe was quantified (Figure 2A-B). The analysis revealed that Sema3C exerts an attractive effect on MGE-derived neurons. 53 ± 1.10 % of the cells were located on the stripes coated with immobilised Sema3C-Fc, whereas just 47 ± 1.10% of the cells were on the interstripes coated with control (n= 96 frames analysed; paired Student’s t-test; ** p ≤ 0.01). The results of the chemotactic stripe assay suggest that Sema3C represents a permissive factor for cortical interneurons.

**Figure 2.**
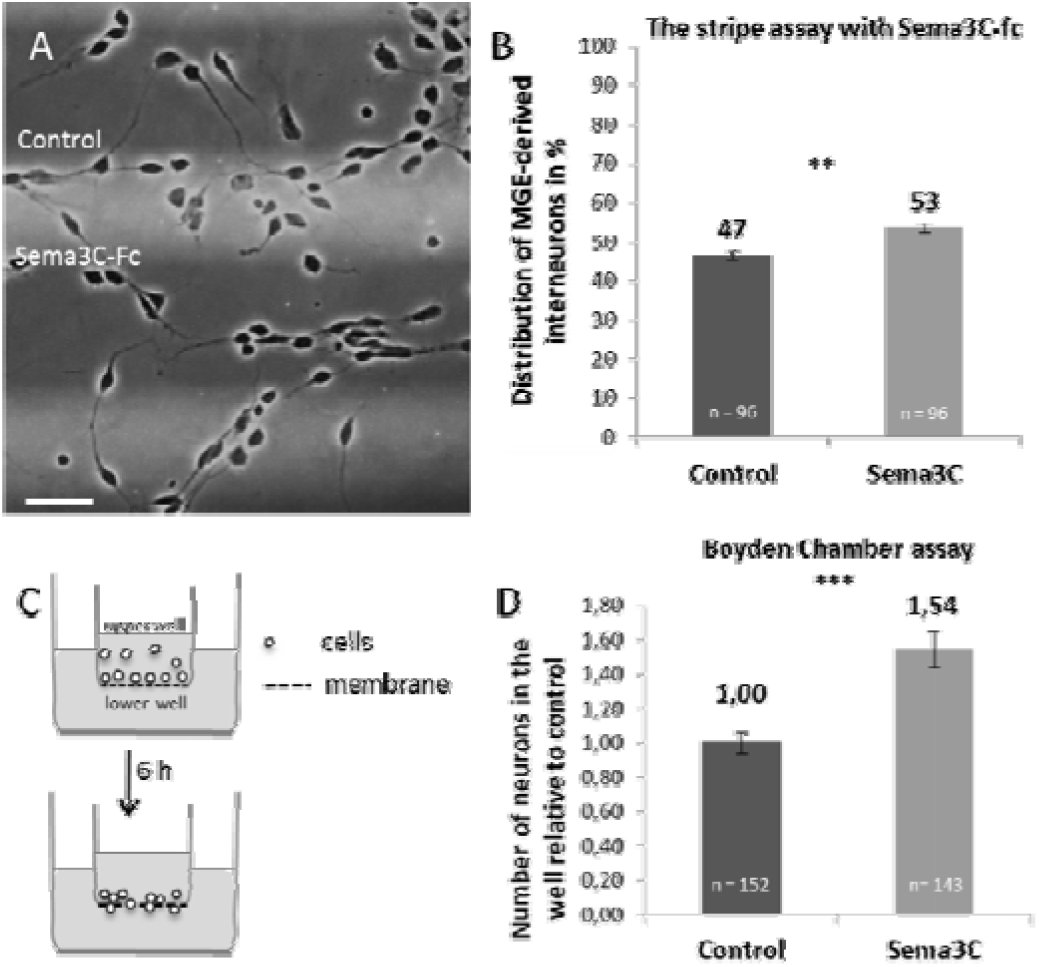
Sema3C-Fc has an attractive effect on MGE-derived interneurons *in vitro*-. (A) Dissociated E14.5 MGE-derived cells cultured on alternating stripes containing only Laminin/PLL (control), or added Alexa488-labeled Sema3C-Fc 50 μg/ml after 2 div. (B) Quantification of the stripe assay after 2 div. Cortical interneurons shows a preference to grow on Sema3C-Fc stripes. Scale bar: 50 μm; n, number of pictures analyzed; Student’s paired t-test: **p < 0.01; error bars are SD. Results are from three independent experiments. MGE, medial ganglionic eminence; Fc, Fragment crystallizable; PLL, poly-L-lysine. (C) Scheme of the Boyden chambers assay. Dissociated E14.5 MGE-derived cells were placed in the upper and the cue tested in the lower compartment of the chamber. Single cells were incubated for 6 h. Migratory cells are able to pass through the pores of the polycarbonate membrane in the presence of an attractive cue. In the case of a neutral cue placed in the lower compartment, cells will pass less through the membrane. (D) The quantification of neurons in the lower well was calculated relative to control, which was set to ‘1’. Values showed that Sema3C-AP gradients exerted attraction on MGE-derived cells. n, number of frames analyzed; Student’s t-test: ***p≤ 0.001, error bars are SEM. Results are from two independent experiments.

**Figure 3.**
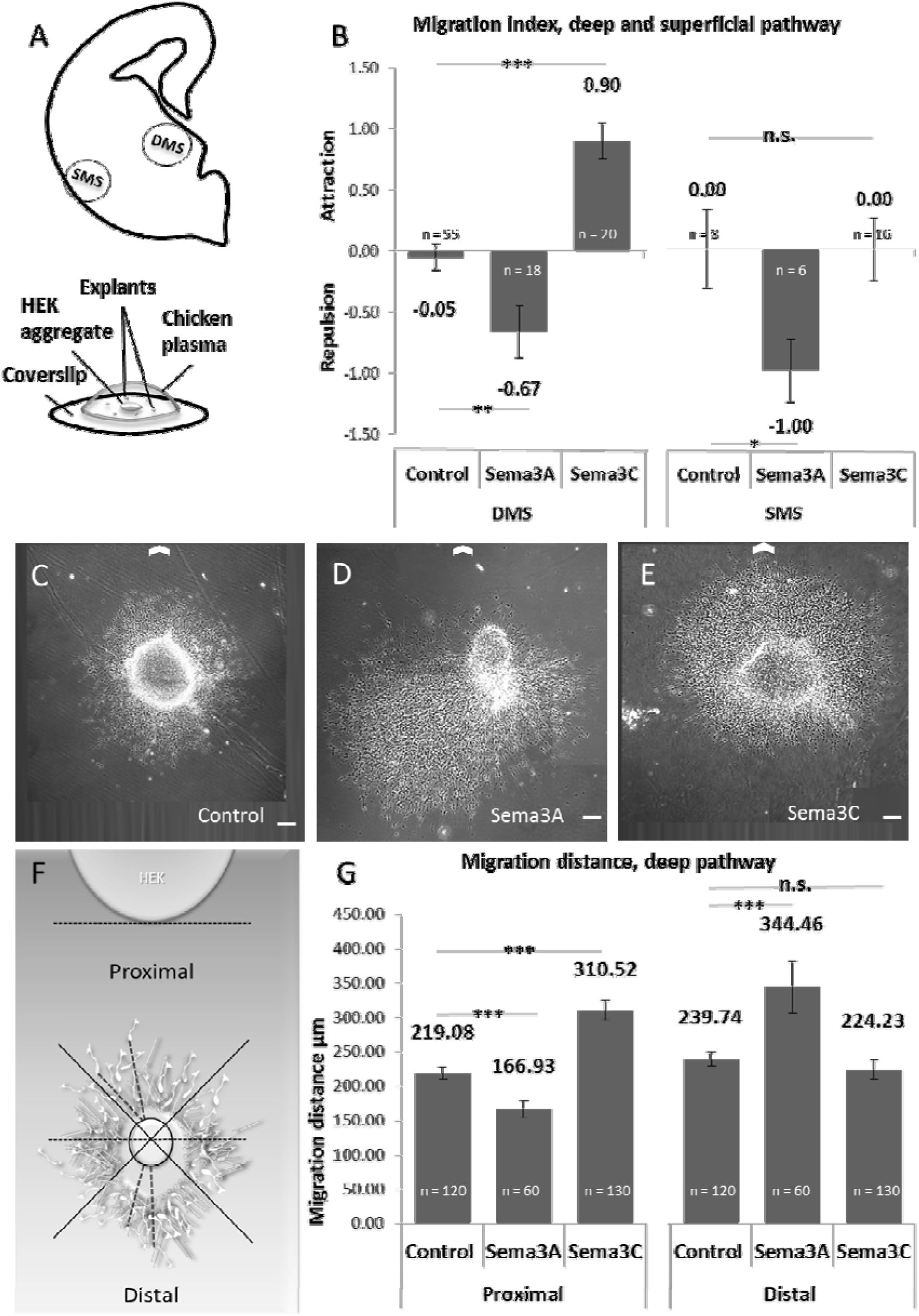
Sema3C and Sema3A gradient-effects on MGE-derived cells from the DMS and SMS. (A) Experimental design, E14.5 coronal slices were used to dissect tissue from the VZ/SVZ or IMZ of MGEs and form explants. One HEK aggregate and 5 to 7 explants were cocultured 2 div in chicken plasma cross-linked with thrombin. (B) Quantification of the guidance index for DMS, using VZ/SVZ, or SMS, using IMZ explants. Bar shows the guidance index values under each coculture condition. Neurons derived from the VZ/SVZ of the MGE are repelled by Sema3A and attracted towards Sema3C. In contrast, neurons derived from the IMZ of the MGE were repelled by Sema3A, whereas Sema3C had no effect on them. n, explants analyzed; Students t-test: ***p < 0.001; error bars are SEM. Results are from at least three independent experiments using VZ/SVZ explants and two independent experiments using IMZ explant. (C, D, E) Response of explants from the VZ/SVZ of the MGE to Sema3A-AP, Sema3C-AP or control gradients in coculture assays. Micrographs of representative MGE explants cocultured with (C) aggregates of non-transfected HEK cells (control), (D) Sema3A-AP transfected HEK cells and (E) Sema3C-AP transfected cells. White arrow represents the position of the HEK aggregates. Scale bars: 100 μm. (F)Design of the experimental quantification. Coculture was performed as described in (A) with VZ/SVZ of MGEs explants only. Analysis was performed after 2 div coculture, and further fixation, for every condition blindly. The initial shape of the MGE was outline with a black line. Proximal and distal areas were set perpendicular to the HEK cell aggregate. In the distal area, light dashed lines illustrate the migration distance of two MGE-derived interneurons, measured from the initial border of the MGE explant to the soma. (G) Bar shows the distance migrated by the MGE neurons on each coculture condition. Neurons migrated shorter distances towards Sema3A increasing gradients, whereas migrated longer distance away from Sema3A decreasing gradients. In contrast, neurons migrated longer distances towards Sema3C increasing gradients, while decreasing Sema3C gradients had no effect on their migration when compared to control conditions. Students t-test: ***p < 0.001; error bars are SEM. Explant used for analysis had a clear initial border. Results are from at least three independent experiments. VZ/SVZ, Ventricular/subventricular zone; IMZ, Intermediate zone; MGE, medial ganglionic eminences. DMS, Deep Migratory Stream ; SMS, Superficial Migratory Stream

In the stripe assay the tested proteins were immobilized, but Class-III Semaphorins represent secreted molecules that form gradients. We next questioned whether soluble Sema3C exerts the same attractive effect on migration interneurons. Thus, we performed another chemotactic assay with diffusible proteins in transwell compartments as recently described [25]. For the Boyden-Chamber assay, we used recombinant Semaphorin3C-AP or control, produced by stable transfected HEK cells that secrete the proteins into the culture medium. The conditioned medium was concentrated and added to the lower compartment of the transwell insert (Figure 2C). The recombinant proteins diffuse through the membrane, thereby forming local gradients. MGE-derived single cells at E14.5 were added to the upper compartment, migrating through the membrane towards their environment of preference. After 6 hours *in vitro* the membrane of the insert was fixed and cells on the side facing the gradient were quantified (Figure 2D). The amount of cells migrating was standardized in relation to the control.

The Boyden chamber revealed an attractive effect of Sema3C on the cultured MGE-derived cells, even stronger than in the stripe assay. Cells migrating through the membrane towards the Sema3C-AP gradient doubled migrating cells under control conditioned medium (Control 1± 0.06; Sema3C-AP 1.5 ± 0.11; Control or Sema3C-AP n = 60 frames analysed; Student’s ttest; *** p ≤ 0.001). The results show the attractive effect of Sema3C on cortical interneurons and further reveal that gradients of Sema3C are more attractive than evenly distributed immobilized protein.

### 3.3. Sema3C gradients exclusively attract interneurons in the Deep Migratory Stream and not in the Superficial Migratory Stream

We further study the effect of free diffusible Sema3C on cortical interneurons following the two different pathways to the neocortex: the deep migratory stream where interneurons mostly expressed Nrp1 and the superficial migratory stream where interneurons expressed mostly Nrp2 receptor, among others [5]. Using the coculture assay we created a three dimensional matrix that closely resembles in vivo conditions. HEK-cell aggregates that either expressed Sema3C-AP, Sema3A-AP or Control were cocultured with MGE-explants from the superficial-or deep-migratory stream of E14.5 embryos. HEK-cells expressing Sema3A-AP were used as positive control.

In this approach, the proteins of interest are constantly secreted by the HEK-cells creating diffusible gradients through the three dimensional gel. To quantify the general effect of the guidance cues, we divided the MGE-explants into four sectors that were directed towards the HEK-cells or away, respectively. It is expected that cells migrate out of the MGE-explants uniformly, moving further towards attractive gradients, whereas chemorepulsive cues direct the migratory cells away from the source of expression. In the coculture assay, the growth of the explant is scored with 0 for even cell migration, 1 or 2 when cells migrate moderately or strong towards the HEK-cell aggregate and −1 or −2 when cells migrate moderately or strong away from the HEK cell aggregate [11]. For each condition, the guidance index was calculated averaging all the score values of the explants.

In the presence of Sema3A-AP, neurons from the DMS- and SMS- explants show an index of −0.67 ± 0.21, and −1 ± 0.04 respectively. These results indicate that Sema3A-AP gradients are repellent for migrating cortical interneurons passing through both migratory streams. As expected, in control conditions cells migrated uniformly out of the DMS- or SMS- explants with guidance indexes of −0.05 ± 0.10 and 0.00 ± 0.30 respectively (Fig. 3B; DMS: Control n = 55, Sema3A-AP n = 18; Student’s t-test, ** p ≤ 0.01; SMS: Control n = 8, Sema3A-AP n = 6; Student’s t-test, * p ≤ 0.05).

In the presence of Sema3C-AP gradients, neurons from the SMS presented a migration index of 0.00 ± 0.25 (Fig. 3B; SMS: Control n = 8, Sema3C-AP n = 16; Student’s t-test, n.s p=1). In contrast, neurons from the DMS migrated more towards the Sema3C-AP gradients obtaining a guidance index of 0.90 ± 0.14 (Fig. 3 B; DMS: Control n = 55, Sema3C-AP n = 20; Student’s ttest, *** p ≤ 0.001). The results confirmed that Sema3C has no effect on cells en route to the cortex through the superficial migratory stream, but attracts neurons migrating through the deep migratory stream.

Next, we further measured the migration distance of the ten farthest neurons that left the MGE-explant from the DMS towards or away from the HEK-cell aggregate. Because it was shown that Sema3C-AP has no effect on interneurons passing through the SMS, only MGEexplants from the DMS were used.

Surprisingly, in the distal area, where the amount of Sema3A-AP protein is more diluted, neurons doubled their migratory distance compared to control conditions, indicated by an average distance of 344.46 ± 37.21 μm. Explants opposed to Sema3A-AP gradients migrated about a third less in the proximal area than under control conditions, with an average distance of 166.93 ± 12.60 μm (Fig. 3G; Control n= 420, Sema3A-AP n = 60 proximal area, Sema3A-AP n = 60 distal area, Student’s t-test, *** p ≤ 0.001). As expected, explants confronted with untransfected HEK cell aggregates migrated out equal distances on each area, with an average value of 229.41 ± 6.73 μm (Fig. 3G; Control n = 420, Control n=210 proximal area vs Control n=210 distal area, Student’s t-test, n.s. p = 0.125).

Under the Sema3C-gradient, in the distal area, where the amount of Sema3C-AP protein is more diluted, the migrated distances were similar to control conditions, with an average value of 224.23 ± 14.50 μm (Fig. 3G; Control n = 420, Sema3C-AP n = 130 distal area; Student’s t-test, n.s. p = 0.721). Neurons migrated further, around a third more than under control conditions in the proximal area, with an average distance of 310.52 ± 14.23 μm (Fig. 3G; Control n = 420, Sema3C n = 130 proximal area; Student’s t-test, *** p ≤ 0.001). Thus, only Sema3C increasing gradients stimulated the movement of MGE-derived inhibitory neurons towards the guidance cue source.

The previous results reinforced the hypothesis that Sema3C is attracting cortical interneurons migrating through the DMS toward the neocortex. Also, the effects are modulated by protein concentration in both Sema3C, and our positive control, Sema3A.

## 4. DISCUSSION

Previous studies have identified molecular mechanisms that guide and regulate interneuron migration. Using different *in vitro* assays, we found a novel role for Sema3C, showing an attractive effect on MGE-derived interneurons migrating through the deep migratory stream.

### 4.1. Semaphorin 3C effects during tangential migration of cortical interneurons

Cortical interneurons derived from the basal telencephalon and migrate long distances to their final position in the developing cortex [5, 6, 10, 11]. Guiding molecules together with cell-intrinsic programs, translated in signalling receptors, assure that cortical interneurons reach their target areas. Several classes of guidance molecules, including Semaphorins, have been implicated in the regulation of interneuron migration creating permissive and nonpermissive environments that are used by cortical interneurons to enter the neocortex. [6–8, 12–17, 28]

Among the Semaphorin family, Sema3A and Sema3F are secreted in the striatum preventing cortical interneurons from entering into this developing structure, thus moving forward to their target areas in the dorsal telencephalon [7, 17, 27, 28].

Previous results showed that Sema3C expression patterns sharply demarcate the pallial/subpallial border and the intracortical pathway of migrating interneurons in the subventricular zone of the neocortex [18, 19]. Sema3C is a secreted protein and may form a gradient that reaches cortical interneurons on their route from the basal telencephalon towards the neocortex. Our study indicated that increasing concentrations of Sema3C attracts MGE-derived cortical interneurons migrating through the deep migratory stream. Also, while using Sema3A as positive control our results showed that a decreasing concentration of Sema3A almost doubled the migration distance of MGE-derived interneurons *in vitro*. Would be of interest to elucidate to what extent this gradient dependent effect is repellent or proliferative as suggested by recent works from Andrew et al., [6].

### 4.2. Neuropilins as Sema3 coreceptors and the overlap of Sema3A and Sema3C opposing cues

The Semaphorin family signals through specific receptor complexes. An interesting feature of Class III Semaphorin is Neuropilins function as a coreceptor. Due to the small cytoplasmic domain, Neuropilins need other molecules, generally Plexins, to transduce their signals across the cell membrane [30].

Sema3A repulsion on migrating cortical interneurons is mediated by Nrp1 coreceptor and Limk2-PlexinA1 [6, 7, 28, 31]. We found in our coculture assay that Sema3A also repels cortical interneurons migrating through the superficial migratory stream, where Nrp2 receptors are expressed instead of Nrp1 or Limk2-PlexinA1. Thus, it remains to be seen which receptor complexes are signalling Sema3A repellent-effects on cortical interneuron migration through the superficial pathway.

In addition, Sema3C and Sema3A gradient might overlap in the basal telencephalon as they overlap in the dorsal telencephalon. The overlapping of opposing cues raises the questions: *How do migrating interneurons following the deep migratory stream translate Sema3A and Sema3C, while signalling through the same coreceptor Nrp1?* Also, *is this overlapping playing a role in the intracortical migration of interneurons, confining them through the SVZ and the MZ of the neocortex?* One possibility is that interneurons use Nrp1 modulated by other molecules when encountering overlapping Sema3A and 3C cues. Hereafter, Sema3A repulsion could override Sema3C attraction in interneurons, as previously reported for cortical axons [18]. A second possibility is that specific Class III Semaphorin-binding desensitises migrating interneurons to other guidance molecules [32, 33]

### 4.3. Attractive guidance cues Nrg1 / Cxcl12

So far, one long and short range molecule described as attractive for a subpopulation of migrating MGE-derived interneurons is Neuroregulin 1 (Nrg1). Nrg1 signals through ErbB4 receptors. Membrane bound Nrg1-CRD (Cysteine Rich Domain) creates a permissive corridor that channels ErbB4 positive MGE-derived interneurons through the lateral ganglionic eminence (LGE), helping to avoid the striatum. On the other hand, secreted Nrg1-Ig (Immunoglobulin-like domain) isoforms strongly attract a subpopulation of MGE-derived interneurons that bear the ErBb4 receptor towards the neocortex [16]. Since ErbB4 is only expressed by a subpopulation of interneurons, the existence of other long range attractive molecules that might guide subpopulation of cortical interneurons towards the neocortex seems likely. Also, Cxcl12 is a diffusible protein modulated by CxCR7 receptors and likely restrained by heparan sulfates [32]. Cxcl12 creates attractive corridors to guide CxCR4-positive MGEderived interneurons in the neocortex and seems to control their sorting and laminar positioning on the cortical layers. Such corridors are in the marginal zone and the subventricular zone of the neocortex [35–40]. The similarity in the attractive role and expression patterns of Sema3C and Cxl12, raises the possibility that Sema3C might also play a role in confining migrating interneurons into specific routes towards the neocortex.

## 5. CONCLUSION

The present study confirms that migrating MGE-derived interneurons are repelled by increasing concentrations of Sema3A. In contrast, we show that Sema3C represents a new permissive cue during brain development, attracting Nrp1-cortical interneurons to the dorsal telencephalon. As a diffusible cue, Sema3C is released and forms gradients and it seems to act as a long-range cue in Nrp1-interneurons migrating through the deep migratory stream towards Sema3C-producing cells in the neocortex.

